# Fungal-bacterial gut microbiota interactions in patients with *Clostridioides difficile* colonisation and infection

**DOI:** 10.1101/2023.07.12.548349

**Authors:** Jannie G.E. Henderickx, Monique J.T. Crobach, Elisabeth M. Terveer, Wiep Klaas Smits, Ed J. Kuijper, Romy D. Zwittink

## Abstract

**Objectives:** The bacterial microbiota is well-recognised for its role in *Clostridioides difficile* colonisation and infection, while fungi and yeasts remain understudied. The aim of this study was to analyse the mycobiota and its interactions with the bacterial microbiota in light of *C. difficile* colonisation and infection.

**Methods:** The mycobiota was profiled by ITS2 sequencing of faecal DNA from infected patients (CDI; n = 29), asymptomatically colonised patients (CDC; n = 38) and hospitalised controls with *C. difficile* negative stool culture (Controls; n = 38). Previously published 16S rRNA gene sequencing data of the same cohort were used additionally for machine learning and fungal-bacterial network analysis.

**Results:** CDI patients were characterised by a significantly higher abundance of *Candida* spp. (MD 0.270 ± 0.089, *P* = 0.002) and *Candida albicans* (MD 0.165 ± 0.082, *P* = 0.023) compared to Controls. Additionally, they were deprived of *Aspergillus* spp. (MD -0.067 ± 0.026, *P* = 0.000) and *Penicillium* spp. (MD -0.118 ± 0.043, *P* = 0.000) compared to CDC patients. Network analysis revealed a positive association between several fungi and bacteria in CDI and CDC, although the analysis did not reveal a direct association between *Clostridioides* spp. and fungi. Furthermore, the microbiota machine learning model outperformed the models based on the mycobiota and the joint microbiota-mycobiota model. The microbiota classifier successfully distinguished CDI from CDC (AUROC = 0.884) and CDI from Controls (AUROC = 0.905). *Blautia* and *Bifidobacterium* were marker genera associated with CDC patients and Controls.

**Conclusions:** The gut mycobiota differs between CDI, CDC, and Controls, and may affect *Clostridioides* spp. through indirect interactions. The identification of bacterial marker genera associated with CDC and Controls warrants further investigation. Although the mycobiota’s predictive value of *C. difficile* status was low, fungal-bacterial interactions might be considered when diagnosing and treating *C. difficile* infection.

## INTRODUCTION

*Clostridioides difficile* is an anaerobic, Gram-positive, spore-forming bacterium and the main causative organism of nosocomial diarrhoea. Upon secretion of toxin A and B, colonic inflammation is induced, giving rise to clinical manifestations that range from mild diarrhoea with abdominal cramping to life-threatening pseudomembranous colitis, toxic megacolon, and death [1].

The disruption of the microbiome due to antibiotic use is a major risk factor for *C. difficile* infection (CDI), as is an age of 65 years and above [1]. The microbiome of CDI patients is characterised by low diversity, decreases in the abundance of *Bacteroides, Prevotella* and *Bifidobacterium* species, while the abundance of *Clostridioides* and *Lactobacillus* species are increased [2,3]. It remains unknown if these microbial changes are associated to risk factors leading to CDI, such as antibiotic use, or due the presence of *C. difficile* in the gut [4]. Faecal microbiota transplantation (FMT) is a highly effective treatment for recurrent CDI (rCDI) that re-establishes a healthy gut microbial community, inhibits the growth of *C. difficile* and prevents recurrence [5–10].

Although *C. difficile* can lead to clinical manifestations (CDI), the organism can also be part of the human gut microbiota without causing symptoms. Such asymptomatic colonisation is estimated to occur between 0% and 15% of the healthy adult population [11]. Carriage of *C. difficile* is potentially a risk factor for transmission to a susceptible population and, because of the bacterium’s opportunistic nature, it may progress to infection under circumstances in which the microbiome is disturbed [12–16], though carriage does not contribute to transmission in a setting with low prevalence of hypervirulent PCR ribotype 027 [17].

Aside from bacteria, fungi are an integral part of the microbiome. Recent research has emphasised the role of fungi in human health [18]. In CDI specifically, the gut mycobiota has been shown to deviate from the mycobiota of both *C. difficile* carriers and from patients with diarrhoea that is not attributable to CDI [19–22]. The mycobiota of CDI patients is characterised by lower biodiversity, an increased ratio of Ascomycota to Basidiomycota and decreased abundance of *Saccharomyces, Cladosporium* and *Aspergillus* [19–21,23]. Additionally, the effectiveness of FMT for treatment of CDI has been positively associated with the genera *Saccharomyces* and *Aspergillus* and negatively associated to dominance of *Candida albicans* [24].

Importantly, interactions between fungi and bacteria have been described in CDI pathophysiology, as *C. difficile* was shown to withstand otherwise toxic aerobic conditions when cultured in presence of *Candida albicans* [25]. Further supporting the importance of fungal-bacterial interactions, it was shown that *C. difficile*-directed antibiotics, including metronidazole and vancomycin, lead to outgrowth and emergence of potentially pathogenic fungi [23]. As such, an intriguing triad of antibiotics, bacteria and fungi exists in *C. difficile* colonisation and infection [26].

Currently data on the gut mycobiota in relation to *C. difficile* carriers and infected patients is limited, and the fungal-bacterial interactions remain unexplored. Moreover, the comparison between *C. difficile* patients and hospitalised controls, who have comparable comorbidities and received antibiotic treatment, is currently missing. Here, we aim to unravel the role of the mycobiota and to broaden our understanding of fungal-bacterial interactions in *C. difficile* colonisation and infection through a comparison with hospitalised, non-colonised controls.

## METHODS

Patients were part of one of the three study groups: patients with *C. difficile* infection (CDI), asymptomatic *C. difficile* colonisation (CDC) and patients without *C. difficile* colonisation (Controls). Microbiological and microbiota analyses have been performed and processed as previously described [26]. The institutional review board of Leiden University Medical Center (LUMC) and the directing board of the Amphia hospital had no objection to the performance of the study. A waiver for informed consent for stool collection of CDI patients was obtained. Faecal samples of CDC patients and Controls were collected under verbal consent; written informed consent was obtained for collection of additional data [27].

In the current study, available faecal samples (Total: n = 105; CDI: n = 29; CDC: n = 38; Controls: n = 38) were used for mycobiota profiling. DNA extraction, quality control, library preparation and ITS2 sequencing were performed according to standard operating procedures of BaseClear B.V. (Leiden, the Netherlands). During DNA extraction and sequencing, negative controls (empty tubes) and positive controls (D6300 ZymoBIOMICS Microbial Community standard and D6305 ZymoBIOMICS Microbial Community DNA standard, Zymo Research, USA) were included. Sequencing was performed on the MiSeq Illumina platform. Raw reads were processed according to the Q2-ITSxpress workflow [28] and classified with the UNITE database [29]. Subsequent pre-processing, quality control and downstream analyses were performed in R (version 4.1.2) [30]. In short, taxonomic profiles and Shannon diversity were generated for CDI, CDC, and Controls. Moreover, differential abundant features on genus and species level were identified with ALDEx2 (version 1.26.0) [31] using centred log-ratio transformed values and default parameters. Machine learning was performed with the SIAMCAT package (version 2.1.3) [32] on 16S rRNA and ITS2 gene amplicon sequencing separately and on the joint data. Fungal-bacterial network analyses were performed with SParse InversE Covariance Estimation for Ecological Association Inference, SpiecEasi (version 1.1.2) [33] combining 16S rRNA and ITS2 amplicon sequencing data. For detailed methods, please see the supplementary material.

## RESULTS

### *C. difficile*-infected patients are enriched with *Candida* spp

To assess differences in the mycobiota of CDI, CDC and Controls, taxonomic profiles of the mycobiota using ITS2 sequencing were generated (**Figure 1**). This revealed the presence of eukaryotic kingdoms including Fungi, Viridaeplantae and Metazoa. (**Figure 1A**). For further analyses, fungal and unassigned kingdom reads were retained, and comprised 468 out of 959 taxa and 68.8% of total reads. The relative abundance of the fungal kingdom was 0.75, 0.60 and 0.57 in CDI, CDC, and Controls, respectively (± 0.32, 0.39 and 0.39 S.D.).

**Figure 1.**
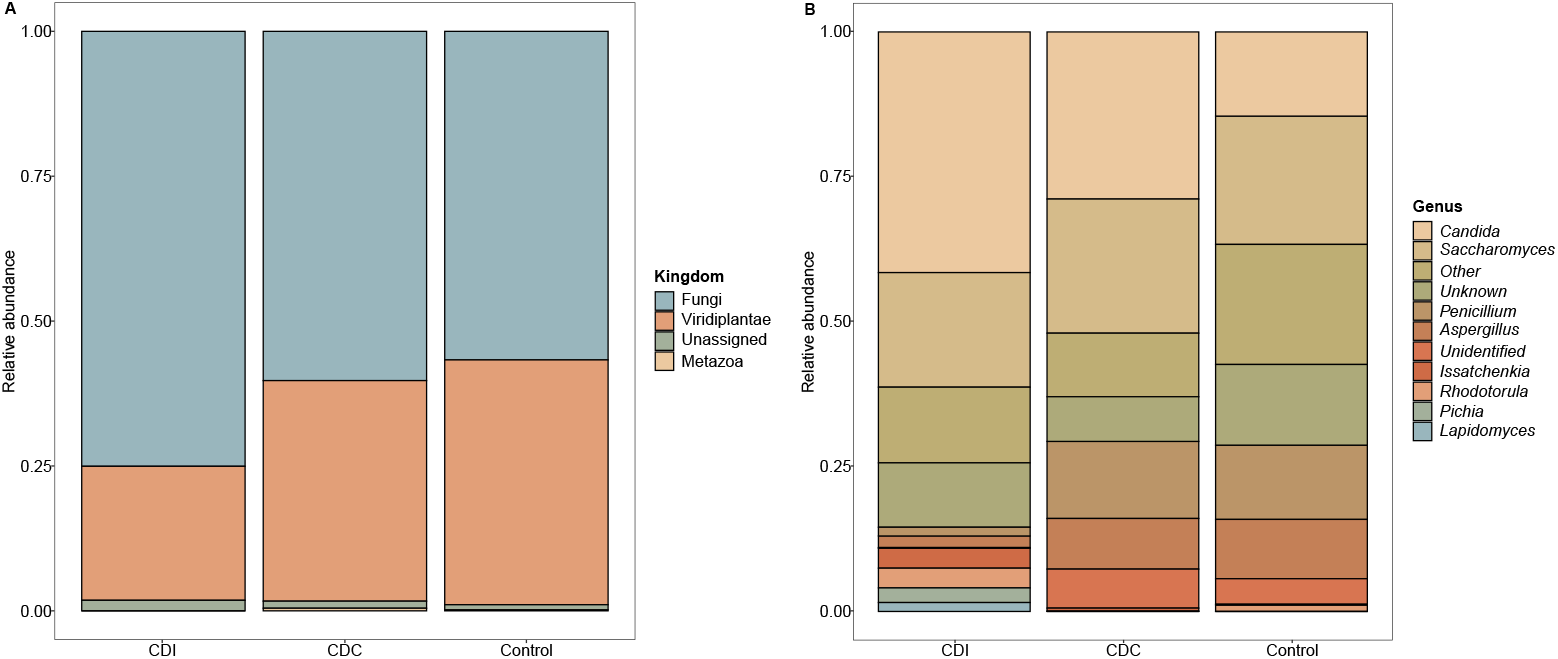
Taxonomic profiles of CDI, CDC and Controls. A: Eukaryotic taxonomic profiles on kingdom level are shown for the three study groups. B: Fungal taxonomic profiles on genus level are shown for the ten most abundant genera. Taxa not belonging to the ten most abundant genera were summarized into ‘Other’.

The most abundant genus in CDI patients was *Candida* spp. (**Figure 1B**), with a significant mean difference (MD) compared to Controls (**Figure 2A**, MD 0.270 ± 0.089, *P* = 0.002, Wilcoxon Rank Sum Test). More specifically, the mean relative abundance of *Candida albicans* was higher in CDI patients compared to Controls (**Figure 2B**, MD 0.165 ± 0.082, *P* = 0.023). This was accompanied by a decrease in *Saccharomyces* spp., with a significant higher *Candida* to *Saccharomyces* ratio in CDI patients compared to Controls (**Supplementary Figure 3**, *P* = 0.024, Wilcoxon Rank Sum Test). The *Saccharomyces* genus was underrepresented in the mock community, which may overestimate the shift in the ratio. Moreover, mean relative abundance of the genera *Aspergillus* and *Penicillium* were significantly lower in CDI patients compared to CDC patients and Controls (**Supplementary Figure 4**).

**Figure 2.**
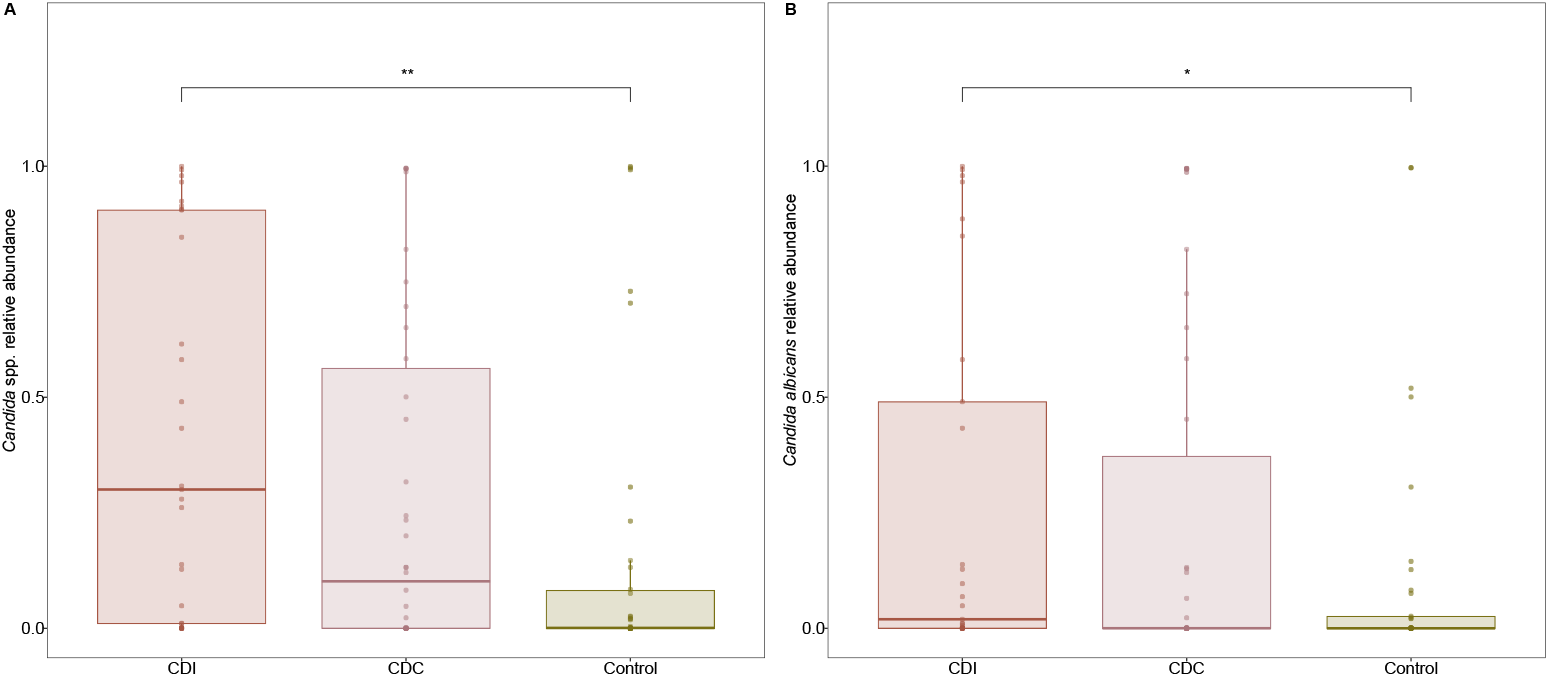
Relative abundance of *Candida* spp. (A) and *Candida albicans* (B) in CDI, CDC and Controls. Individual data points are displayed as circles. **P ≤ 0.01, *P ≤ 0.05 (Wilcoxon Rank Sum Test).

Differential abundance testing confirmed the significant increases in CDI patients of *Candida* spp. (*P*_*unadj*_ = 0.003) and *Candida albicans* (*P*_*unadj*_ = 0.027) compared to the Controls, as well as the significant deprivation of *Aspergillus* (*P*_*adj*_ = 0.044) and *Penicillium* species (*P*_*unadj*_ = 0.002) compared to CDC patients. No differential abundant features were identified between CDC patients and Controls.

Besides differences in the mycobiota composition, the median Shannon diversity increased from CDI patients to CDC patients to Controls, though differences between groups were not significant (**Supplementary Figure 2**, *P* = 0.167, Kruskal-Wallis test).

Together, these results indicate that CDI patients were most deviant from CDC patients and Controls regarding the taxonomic composition and the diversity of the mycobiota, even though majority of the patients from each group had received antibiotic treatment up to three months prior to sample collection [27].

### The microbiota can discriminate *C. difficile* infection from colonisation

Machine learning was used to investigate if models of the microbiota, mycobiota and joint data could discriminate between CDI, CDC, and Controls. With regard to the Controls, antibiotic-treated Controls were selected to exclude bias introduced by antibiotic treatment. The performance of the models was evaluated based on the Area Under the Receiver Operating Characteristic (AUROC), indicating the probability to classify a randomly selected patient to the correct study group.

First, the bacterial microbiota, mycobiota and joint models were used to assess their performance in discriminating between CDI and CDC patients, as well as between CDI patients and Controls. Comparing CDI and CDC patients, the AUROC of the mycobiota (0.627, 95% CI: 0.491 - 0.764) was significantly lower compared to that of the bacterial microbiota (0.884, 95% CI: 0.808 - 0.960) (**Supplementary Figure 5A**). Similar results were obtained when CDI patients were compared to Controls (AUROC mycobiota: 0.632, 95% CI: 0.469 - 0.795, AUROC microbiota: 0.905, 95% CI 0.816 - 0.991) (**Supplementary Figure 5B**). Moreover, the joint model did not significantly differ from the bacterial microbiota models when comparing CDI to CDC patients (0.865, 95% CI: 0.782 - 0.948) and CDI to antibiotic-treated Controls (0.897, 95% CI: 0.810 - 0.984) (**Supplementary Figure 5A and B**). Overall, these results indicate that the bacterial fraction of the microbiota is most successful in discriminating between CDI and CDC patients and between CDI patients and Controls.

The two microbiota models with an AUROC above 0.800 were selected to further assess the bacterial OTUs that were collectively characteristic for CDI and CDC patients (**Figure 3**), and for CDI patients and Controls (**Supplementary Figure 6**). In both cases, *Clostridioides* was identified as distinct taxon in CDI patients. The nucleotide sequence of this *Clostridioides* OTU (OTU 1648581238) demonstrated a 100% sequence identity with *Clostridioides difficile*. Compared to CDC, CDI patients were additionally characterised by the genera *Lactobacillus, Haemophilus, Bacteroides* and *Enterococcus* (**Figure 3B**). Interestingly, CDC and Controls shared OTUs from the *Bifidobacterium* and *Blautia* genera. The nucleotide sequences belonging to the identified *Bifidobacterium* OTUs resulted in 100% sequence identity with species *Bifidobacterium breve* and *Bifidobacterium longum* with subspecies *longum, infantis* and *siullum*, while those belonging to the identified *Blautia* OTUs resulted in 100% sequence identity with *Blautia wexlerae, Blautia luti* and *Blautia massiliensis*. In the CDC patients, *Subdoligranulum* was additionally part of the microbial signature (**Figure 3B**), while *Collinsella* was associated with the signature in the Controls (**Supplementary Figure 6B**). Together, these results demonstrate that CDC patients could not be successfully distinguished from Controls based on machine learning with bacterial data. In comparison to CDI, *Bifidobacterium* and *Blautia* were bacterial marker genera for asymptomatic *C. difficile* carriage and non-carriage.

**Figure 3.**
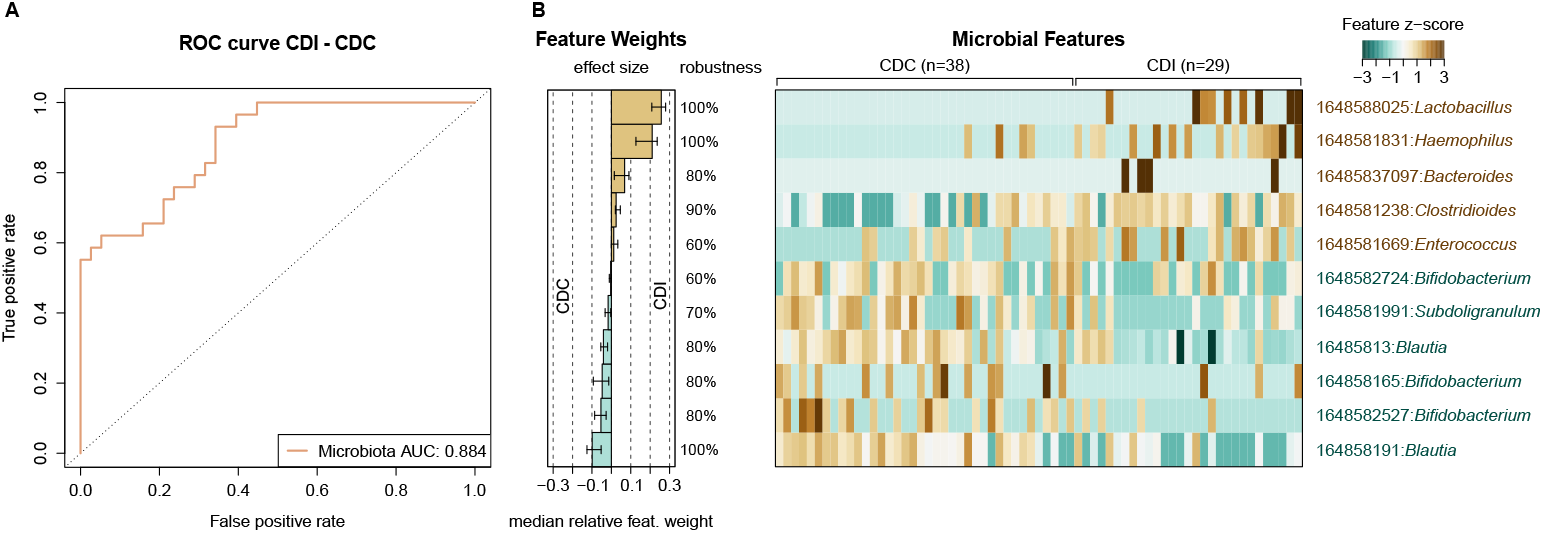
Signature of gut bacterial OTUs associated to CDI and CDC. A: Cross-validation accuracy of the microbiota classifier is shown in the Receiver Operating Characteristic (ROC) curve with the Area Under the Curve (AUC). B: Median relative feature weights of the selected gut bacterial OTUs show the contribution of each marker OTU to the classification. Robustness of the selected features indicates the fraction of models containing the specific feature. The normalised values of the selected features across CDI and CDC is shown in the heatmap.

### Fungi and bacteria interact in *C. difficile* infected and colonised patients

Fungal-bacterial network analysis using SParse InversE Covariance Estimation for Ecological Association Inference (SpiecEasi) was performed for the CDI and CDC patients (**Figure 4**), and for the Controls (**Supplementary Figure 6**). The outcomes of the network analyses were used in combination with the machine learning results to infer potential associations. Interactions were considered for the three most highly abundant fungi of the group that showed interactions with bacteria, as well as for the identified bacterial marker genera (**Figure 3** and **Supplementary Figure 6**). In CDI and CDC patients, bacteria and fungi were only positively associated with each other.

**Figure 4.**
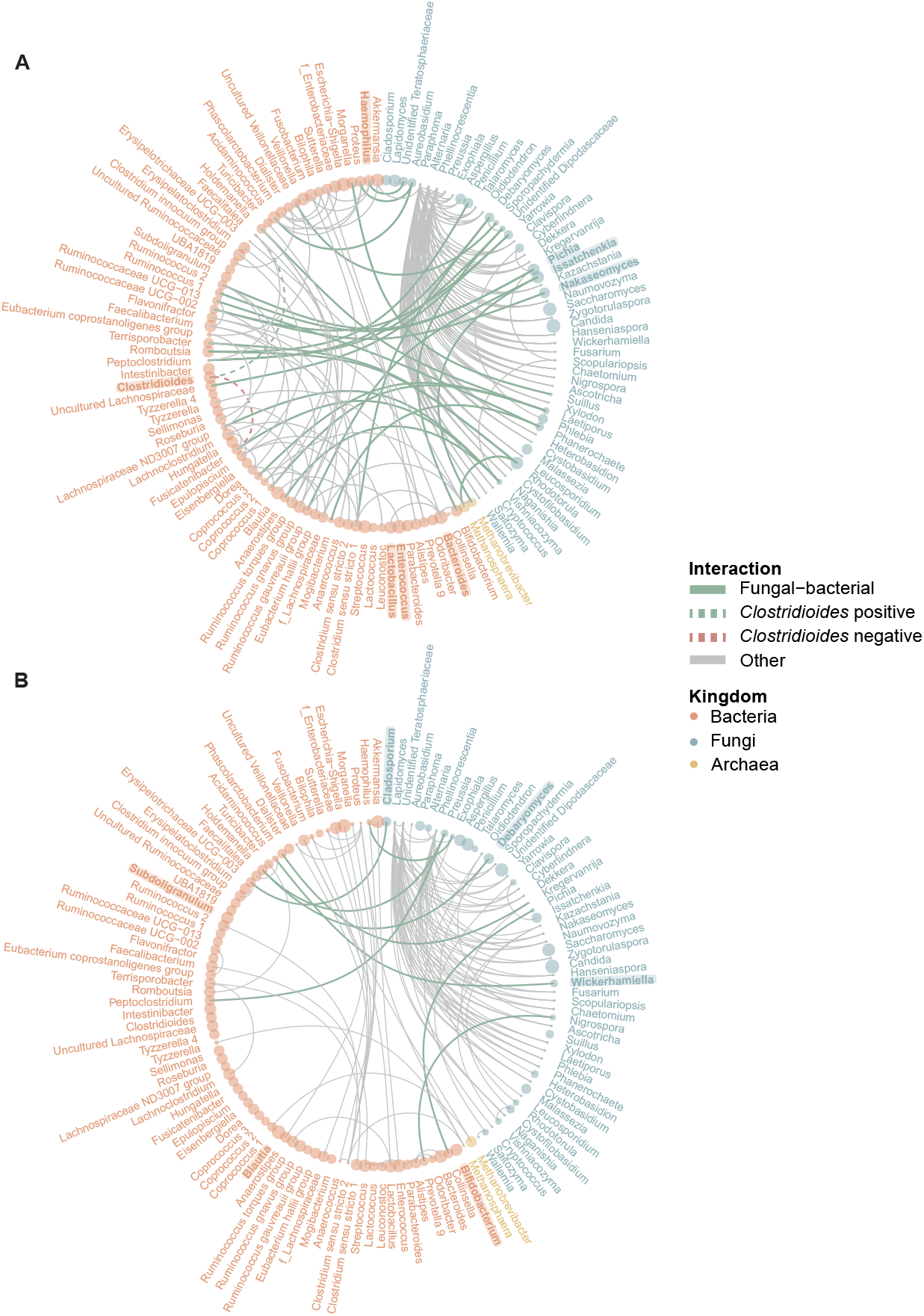
Fungal-bacterial networks in CDI (A) and CDC (B). Interactions between fungi and bacteria are highlighted in green, whilst other interactions are displayed in grey. Kingdoms are represented by colour; mean relative abundance of the genus are represented by point size. Bacterial taxa in bold are part of the microbial signature as identified by machine learning. Fungal taxa in bold are part of the three most abundant fungi in the respective study group. A: In CDI patients, interactions with the *Clostridioides* genus are additionally displayed.

In CDI patients, twenty-nine fungal-bacterial associations were observed (**Figure 4A**). Abundant fungi that positively associated with bacteria included interactions between *Pichia* and *Ruminococcus* 2; *Nakaseomyces* and *Ruminococcaceae* UCG-002; and between *Issatchenkia* and *Ruminococcus gauvreauii* group. Interestingly, no associations were observed for the *Candida* genus. From the bacterial perspective, fungal-bacterial associations were observed for the bacterial marker genera associated to CDI (**Figure 3**). *Haemophilus* positively associated with an unidentified species of the *Teratosphaeriaceae* genus. Yet, the *Clostridioides* genus did not associate to fungi, but did show a negative association to *Fusicatenibacter*. Additionally, no fungal-bacterial associations were observed for the CDI bacterial marker genera *Bacteroides, Bifidobacterium* and *Lactobacillus*, although *Bacteroides* did negatively associate with *Bifidobacterium* and *Lactobacillus* (**Supplementary Figure 7A**).

Nine fungal-bacterial associations were observed in CDC patients (**Figure 4B**). Associations between the abundant fungi and bacteria included interactions between *Debaryomyces* and *Holdemanella*; *Cladosporium* and *Erysipelotrichaceae* UCG-003; and *Wickerhamiella* and *Acidaminococcus*. Similar to CDI, *Candida* spp. did not show any associations. From the bacterial viewpoint, no interactions were observed for the CDC bacterial marker genera (**Supplementary Figure 7B**).

In the Controls who received antibiotics, *Collinsella* and *Bifidobacterium* were few of the bacterial marker genera (**Supplementary Figure 6**). *Collinsella* abundance positively associated with *Anaerostipes*, while *Bifidobacterium* negatively associated with *Aspergillus* (**Supplementary Figure 7C**).

## DISCUSSION

Here, we show distinct fungal profiles in CDI compared to CDC and Controls. We also describe the predictive value of the bacterial microbiota and fungal-bacterial interactions during *C. difficile* colonisation and infection. Mycobiota composition of CDI patients differs from CDC and Controls. Corroborating with previous findings [20,21,24], we report an increase in the ratio between *Candida* spp. and *Saccharomyces* spp., and a deprivation of *Aspergillus* spp. and *Penicillium* spp. in CDI. The enrichment of *Candida* spp. in CDI suggests a decrease in fungal diversity, which could not be confirmed within this study. Current findings on *Candida* spp. enrichment in CDI are inconclusive [19]. This may be explained by different inclusion criteria on antibiotic treatment, which is commonly associated to overgrowth of *Candida* species [34,35]. Indeed, in the study of Stewart *et al*., patients were ineligible whenever *C. difficile*-directed antibiotics were started before stool sample testing [19]. Conversely, 97.6% of CDI and 73.2% CDC participants did receive antibiotic treatment in the last three months prior to faecal sample collection within our study [27]. When we compared CDI patients (n = 29) to Controls who received antibiotics (n = 17), mean *Candida* spp. relative abundance was still significantly higher in CDI patients (*P* = 0.024, data not shown). This suggests *Candida* spp. could be involved in the pathogenesis of CDI rather than being an effect of antibiotic treatment *per se*. Specifically *Candida albicans* could be involved in the pathogenesis of CDI by driving Th17-mediated immune responses and by disrupting the gut microbiome [18]. Additionally, *C. albicans* allows *C. difficile* to grow by reducing oxygen [25]. CDI patients were additionally deprived of *Aspergillus* and *Penicillium. Aspergillus* has been related to healthy status previously [20,21,24] and therefore highlights its potentially beneficial role against CDI. Hypothetically, *Aspergillus* could interact with the immune system by affecting cytokine levels, as this genus was previously negatively correlated to IL-4 and TNF-α serum immune levels in a non-CDI group [20]. Moreover, the significant reduction of the *Candida* to *Saccharomyces* in Controls might point toward a protective effect, as the fungal probiotic *Saccharomyces boulardii* has been associated with a reduced risk of *C. difficile*-associated diarrhoea [36,37]. Moreover, its supplementation during antibiotics prevented development of CDI and reduced rCDI. Yet, evidence is insufficient for clinical application of *S. boulardii as* probiotic [38].

As opposed to CDI patients, CDC patients and Controls were more alike in their bacterial and fungal microbiome. In fact, CDC patients could not be successfully distinguished from Controls based on machine learning with bacterial data alone or when combined with fungal data. *Bifidobacterium* spp. and *Blautia* spp. were bacterial marker genera associated with both CDC and Controls. In line with these findings, other studies have reported depletions of *Bifidobacterium* and *Blautia* in CDI patients compared to non-CDI patients [39] and were associated to less severe CDI-associated outcomes [40]. Among the *Bifidobacterium* OTUs, *Bifidobacterium longum* subsp. *longum* has been identified as a predictor for negative *C. difficile* status [41]. Both *Bifidobacterium* and *Blautia* include species that are known to be involved to bile acid metabolism and short chain fatty acid production [42,43]. In most of the bifidobacterial marker OTUs, bacterial bile salt hydrolase (BSH) genes have been identified. Such mechanisms could contribute to colonisation resistance against *C. difficile* [9,44–46].

The combination of machine learning and network analyses allowed to infer potentially important fungal-bacterial interactions in CDI and CDC patients. Interestingly, *Clostridioides* abundance was not associated with fungi in CDI and CDC in these analyses, while it did identify the previously reported negative association with the Gram-positive *Fusicatenibacter* genus in CDI [27,47]. Moreover, the CDI-associated genus *Bacteroides* showed a negative association with *Bifidobacterium*. As *Bifidobacterium* was a bacterial marker genus associated with CDC patients and Controls, increased relative abundance of *Bacteroides* could have a negative impact on beneficial bacterial-fungal interactions such as those between *Bifidobacterium* and *Aspergillus* as observed in antibiotic-treated Controls. Although explorative, the fungal-bacterial network analyses described herein gives insight into differential interactions that occur during *C. difficile* colonisation and infection.

FMT has proven highly effective for treatment of rCDI, in which the restoration of the bacterial microbiota undoubtedly contributes to successful eradication of CDI symptoms [9]. Emerging data suggest similar roles for the mycobiota. Increased fungal engraftment rates were observed in rCDI patients who responded to FMT compared to patients who did not respond to FMT [24]. More specifically, the genera *Aspergillus, Penicillium* and *Saccharomyces* were significantly higher in responders post-FMT, while a dominance of *Candida* and *C. albicans* was observed in non-responders post-FMT [24]. Additionally, the efficacy of FMT in clearing *C. difficile* infection could be restored when *C. albicans*-colonised mice were treated with the antifungal agent fluconazole before FMT [24]. Within this study, CDI patients were depleted of *Aspergillus, Penicillium* and *Saccharomyces* species while relative abundance of *Candida* spp. and *C. albicans* was increased. Although the mycobiota did not have a predictive role in distinguishing CDI, CDC and Controls, gut fungi might play a role in FMT effectiveness by interacting with the bacterial community of the gut microbiota.

To the best of our knowledge, this study is the first to combine microbiota and mycobiota data in a well-defined cohort of *C. difficile* infected patients, *C. difficile* carriers, and hospitalised non-colonised controls. It was noted in our previous work that patient groups may not be comparable with regard to solid organ transplants, previous hospitalisation, immunosuppressant use and chemotherapy, as majority of diagnosed CDI patients were recruited at the university-affiliated hospital (LUMC) whereas CDC patients and Controls were more evenly recruited at a general hospital (Amphia hospital) and the LUMC [27]. Several of those clinical variables affect microbiota composition [48,49], yet its effect on the mycobiota remain to be elucidated. In a subgroup analysis on LUMC patients, results remained similar (data not shown). While the relative abundance of *Candida* spp. and *Candida albicans* was not significantly higher in CDI compared to all Controls, the mean ratio between *Candida* spp. and *Saccharomyces* spp. remained significantly higher in CDI compared to antibiotic-treated Controls (*P* = 0.018). Moreover, CDI patients were on average depleted of *Aspergillus* spp. (*P* = 0.003) and *Penicillium* spp. (*P* = 0.015) compared to all Controls and to CDC patients, respectively. The relative abundance of specific fungal genera, such as *Candida* spp., might thus be affected by aforementioned clinical variables. Moreover, 16S rRNA gene sequencing and ITS2 sequencing were performed separately. Future studies should rely on shotgun metagenomic sequencing to characterise additional kingdoms as well as genes to perform functional characterisation of the microbiota and mycobiota. The characterisation of additional kingdoms could be especially relevant to investigate the role of the relatively understudied Archaea and phages, which gained increasing attention with regard to FMT efficacy in CDI patients [50].

In conclusion, we have shown that the gut mycobiota differs between *C. difficile* infection, asymptomatic carriage, and non-carriage, as well as their potential fungal-bacterial interactions, which may affect *Clostridioides* spp. The identification of bacterial marker genera associated with carriage and non-carriage warrants further investigation. Although the mycobiota’s predictive value for *C. difficile* status was low, fungal-bacterial interactions might be relevant for microbiota-based strategies in treatment of (recurrent) *C. difficile* infections.

## TRANSPARENCY DECLARATION

## Acknowledgments

We are indebted to all patients who participated in this study. We would like to thank Melanie Srodzinski, Inge van Duijn, Michelle de Raaf and René Vermaire for their help in obtaining faecal samples. Additionally, we would like to thank the authors of the microbiota-based paper of this study and the persons acknowledged therein [27]. Preliminary results of the current study were presented (poster) at the annual meeting of the Royal Dutch Society of Microbiology (KNVM) (Arnhem, the Netherlands, 4-5 April 2023).

## Access to data

The raw ITS (mycobiota) and 16S rRNA (microbiota) gene amplicon sequencing data have been deposited at the European Nucleotide Archive (ENA) at EMBL-EBI under accession number PRJEB61263 and PRJEB30586, respectively.

## Contribution

Made substantial contributions to conception and design of the study: Smits WK, Kuijper EJ, Zwittink RD

Obtained funding: Kuijper EJ, Zwittink RD

Performed data acquisition and supervised the clinical study: Crobach MJT, Terveer EM, Kuijper EJ

Performed bioinformatic analyses: Henderickx JGE

Involved in result interpretation and manuscript preparation: Henderickx JGE, Smits WK, Kuijper EJ

Critically revised the manuscript: Crobach MJT, Terveer EM, Smits WK, Kuijper EJ, Zwittink RD

Reviewed and agreed to the published version of the manuscript: All authors

