## Supplementary Material for "Fungal-bacterial gut microbiota interactions in patients with *Clostridioides difficile* colonisation and infection"

### SUPPLEMENTARY METHODS

#### Subjects and sample collection

Patients were part of one of the three study groups: patients with *C. difficile* infection (CDI), asymptomatic *C. difficile* colonisation (CDC) and patients without *C. difficile* colonisation (Controls). The majority of the patients from each group had received antibiotic treatment up to three months prior to sample collection [1]. CDC and Control samples derived from a multicentre prospective, hospital-based, case-control study that was performed in the Netherlands, as described earlier [1]. The aim of that study was to assess the prevalence of *C. difficile* colonisation at hospital admission and risk of subsequent infection or onward transmission [2]. A subset of samples from this study from two of the participating hospitals (Leiden University Medical Centre (LUMC), a tertiary academic hospital; Amphia hospital, a large general hospital) was included in our analysis. Faeces was collected within 72 hours of hospital admission and cultured for *C. difficile*. Suspicious colonies were tested by glutamate dehydrogenase gene-PCR to confirm *C. difficile* presence and a multiplexPCR on confirmed *C. difficile* isolates was performed for TdcA, TcdB and binary toxin genes. Patients with a positive *C. difficile* culture, but no diagnosis of CDI within the first 72 hours of admission, were considered *C. difficile* colonised (CDC). Patients with a negative stool culture at hospital admission were included as Controls. Samples of the CDI group derived from patients hospitalised at the LUMC and diagnosed with CDI during the study period. CDI diagnosis was based on strict clinical criteria in combination with laboratory CDI testing (including detecting free *C. difficile* toxins and culturing). *C. difficile* isolates from CDI and CDC were additionally ribotyped with capillary PCR [3].

The institutional review board of Leiden University Medical Center (LUMC) and the directing board of the Amphia hospital had no objection to the performance of the study. A waiver for informed consent for stool collection of CDI patients was obtained. Stool samples of CDC and Controls were collected under verbal consent; written informed consent was obtained for collection of additional data.

#### Microbiota analysis

Microbiological and microbiota analyses have been performed and processed previously [1]. ITS2 sequencing was performed on available samples (Total: n = 105; CDI: n = 29; CDC: n = 38; Controls: n = 38) and corresponding samples were selected from the available and processed 16S rRNA sequencing data.

#### Mycobiota analysis

##### *DNA extraction, library preparation and sequencing*

Based on sample availability, mycobiota profiling was performed on 105 out of the 125 faecal samples on which 16S rRNA gene sequencing was performed [1]. DNA extraction, quality control, library preparation and ITS2 sequencing were performed according to standard operating procedures of BaseClear B.V. (Leiden, the Netherlands). DNA was extracted by a combination of chemical and mechanical lysis, in which two empty tubes were included as negative controls and two cellular microbial standards as positive controls (D6300 ZymoBIOMICS Microbial Community standard, Zymo Research, USA). The microbial standard contains ten microbial strains, of which two are fungi: *Saccharomyces cerevisiae* and *Cryptococcus neoformans*. After DNA extraction, concentrations were measured with Quant-iT dsDNA Assay kits (high sensitivity (Q33120) and broad range (Q33130), Invitrogen, USA) and DNA integrity was checked on agarose gel. Subsequently, the Internal Transcribed Spacer (ITS) 2 region was PCR-amplified with primers ITS3 (Forward: 5'-GCATCGATGAAGAACGCAGC-3') and ITS4 (Reverse: 5'-TCCTCCGCTTATTGATATGC-3'), resulting in a ~490 bp amplicon complemented with standard Illumina adapters. Unique Index Primers were attached to amplicons in each sample with a second PCR cycle. PCR products were purified using Agencourt AMPure XP (Becker Coulter) and DNA concentration was measured by fluorometric analysis (Quant-it dsDNA Assay Broad Range kit, Invitrogen). Next, PCR amplicons were equimolarly pooled. Libraries were prepared with Nextera XT Index Kit v2 (FC-131-2004, Illumina, USA), checked on concentration with Quant-iT™ dsDNA Assay High Sensitivity kit and on size with Agilent DNA 1000 (Agilent, USA). After quality control of the library, ITS2 amplicon sequencing was performed on the MiSeq platform (Illumina, USA) with 300 bp paired-end reads. To control for the sequencing process, a microbial DNA standard was included (D6305 ZymoBIOMICS Microbial Community DNA standard, Zymo Research USA).

##### *Bioinformatic analyses: taxonomic assignment*

Raw reads were processed according to the Q2-ITSxpress workflow [4]. Raw reads without barcodes and primers were imported in Qiime2 (version 2022-2) [5]. Subsequently, the conserved regions around the ITS gene were trimmed with ITSxpress, which has been shown to improve accuracy of taxonomic classification [6]. The sequence variants were then identified in the unmerged, trimmed sequences with Dada2 [7]. Next, the Qiime classifier was trained using the UNITE database (version 8.3, all eukaryotes)

[8]. Fungal ITS classifiers were trained on the UNITE database on full reference sequences. Subsequently, sequence variants were classified with the trained classifier.

##### *Bioinformatic analyses: pre-processing data*

Data were imported in R (version 4.1.2) [9] with the *Qiime2R* package (version 0.99.6) [10]. In 105 samples and 6 controls, 959 taxa were identified with 1,721,901 reads in total. The average number of reads were 15,513, with a minimum of 255 and a maximum of 59,126. The sum of reads of three samples were outliers with 21, 255, and 380 reads. These samples were retained since one of those samples was a negative control (NC1) and two samples derived from CDI patients who received antibiotics in the past three months. ASVs were filtered on kingdom, thereby excluding 491 non-fungal taxa consisting of Viridiplantae and Metazoa, while unassigned reads were retained. Filtering resulted in 1,184,306 reads in total, with a mean of 10,669 reads, a minimum of 21 and a maximum of 54,775.

##### *Bioinformatic analyses: data quality control*

Positive controls were included during DNA extraction (PCE) and sequencing (PCS), as well as negative controls for DNA extraction (NC) (**Supplementary Figure 1**). The total number of reads from the filtered ASV table were visualised with *ggplot2* (version 3.3.6) [11]. The mean total reads of positive (22.915 ± 2.686) and negative (748 ± 1.029) DNA extraction controls were not significantly different (**Supplementary Figure 1A**, $P$ = 0.33, Wilcoxon Rank Sum Test). One of the negative controls, NC2, contained *Malassezia globosa* that is often identified on human skin [12], as well as *Cyberlindnera jadinii* (teleomorph of *Candida utilis*) that has been isolated from distinct environments [13] (**Supplementary Figure 1B**).

Additionally, taxonomic compositions of positive controls were compared to the theoretical mock community. The two genera of the mock community — *Cryptococcus* and *Saccharomyces* — are present in a 1:1 ratio in the mock community, indicating a mean overrepresentation (1.48-fold) of *Cryptococcus* spp. and a mean underrepresentation (0.52-fold) of *Saccharomyces* spp. in the DNA and sequencing positive controls (**Supplementary Figure 1C**). Correlation coefficients between the positive controls of DNA extraction ($\rho$= 1.00, $P \leq 0.001$) and between positive controls of sequencing ($\rho$= 0.99, $P \leq 0.001$) indicate high reproducibility between batches.

*Bioinformatic analyses: downstream analyses*

Pre-processed reads were used for downstream analyses in R (version 4.1.2) [9].

Number of reads and relative abundances from the ASV table were used to generate composition plots with *microbiome* (version 1.16.0) [14] and *ggplot2* (version 3.3.6) [11].

Shannon diversity was computed on ASV level with *microbiome*, after reads had been rarified to the minimum sum of reads per sample with *phyloseq* (version 1.38.0) [15]. Overall significant differences were computed with a Kruskal-Wallis test from *stats* (version 4.1.2) [9]. *P*-values below 0.05 were considered statistically significant.

Differential abundant features were identified with *ALDEx2* (version 1.26.0) [16] using centred log-ratio transformed values and default parameters, comparing CDI with Controls and CDI with CDC on genus and species level. The Wilcoxon Rank Sum Test of *ALDEx2* was used for significance testing with Benjamini-Hochberg corrected *P*-values. *P*-values below 0.05 were considered statistically significant. Relative abundance of the differential abundant features were visualised for the three study groups and means were compared with the Wilcoxon Rank Sum Test of *ggpubr* (version 0.4.0) [17].

Machine learning was performed on OUT and ASV level with the SIAMCAT package (version 2.1.3) [18] on 16S rRNA and ITS2 gene amplicon sequencing separately as well as on the joint data. The CDI group was compared to the CDC group and to Controls who received antibiotic treatment. Unsupervised feature filtering was performed with default parameters. Data was normalised with log.unit normalisation and normalisation parameters log.n0 = 1e-06, n.p = 2 and norm.margin = 1. A twice-repeated 5-fold cross-validation scheme was generated and the model was trained using the lasso method.

Fungal-bacterial network analyses were performed with SParse InversE Covariance Estimation for Ecological Association Inference, *SpiecEasi* (version 1.1.2) [19] combining 16S rRNA and ITS2 amplicon sequencing data with *multi.spiec.easi*. Prior to the network analyses, taxa were aggregated to genus level and pruned at 100 reads using *microbiome* and *phyloseq*. Network analyses for each study group were performed using meinshausen-buhlmann's neighborhood selection with lambda.min.ratio = 0.01, nlambda = 20 and rep.num= 99. Networks were visualised with *ggraph* (version 2.0.5) [20].

#### SUPPLEMENTARY FIGURES

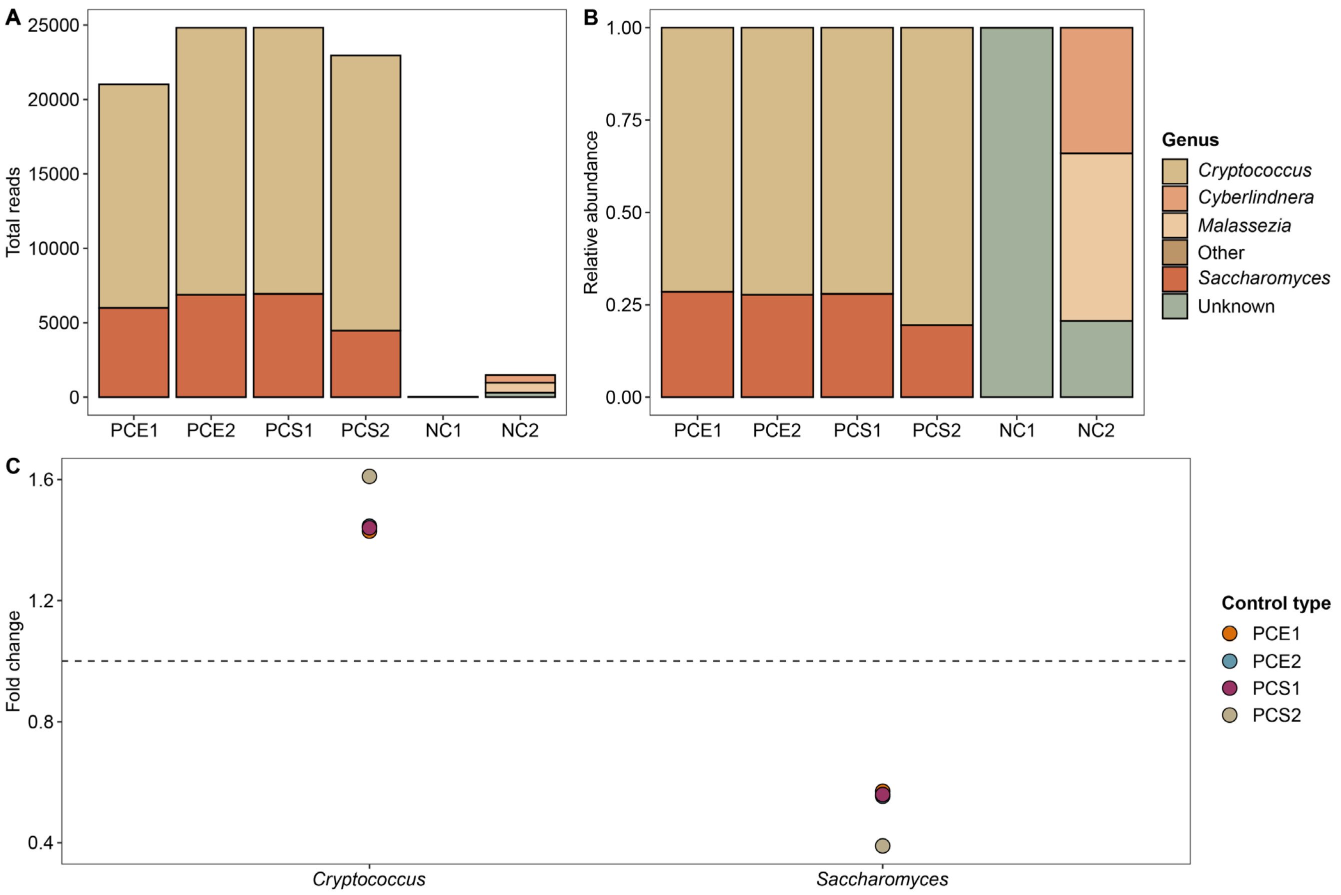


**Supplementary Figure 1.** Positive and negative DNA extraction controls (PCE and NC) and positive sequencing controls (PCS). Fungal taxonomic composition of the five most abundant genera shown as total reads (A) and as relative abundance (B). Taxa not belonging to the five most abundant genera were summarised into "Other". C: Fold change in the relative abundance of the two fungal mock genera compared to the theoretical relative abundance in the mock community (horizontal line).

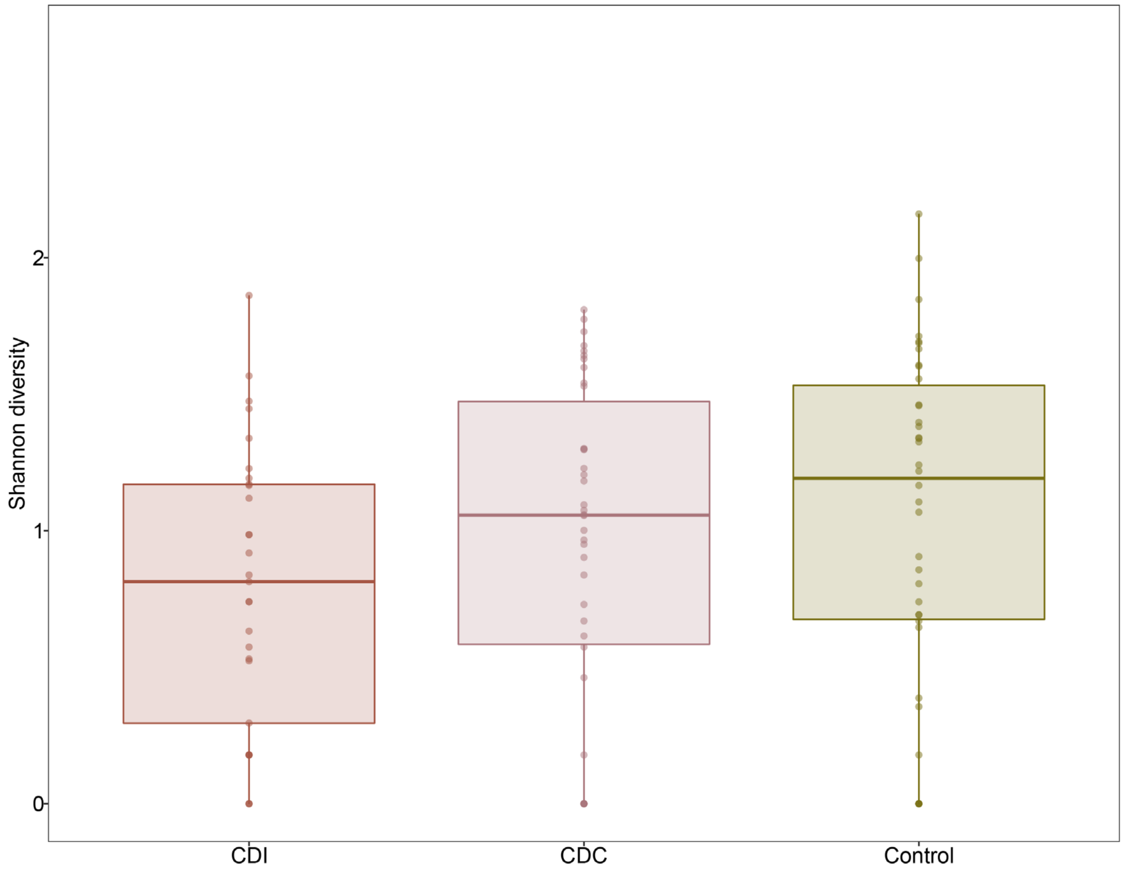


**Supplementary Figure 2.** Shannon diversity of the mycobiota in patients with *C. difficile* infection (CDI), asymptomatic *C. difficile* colonised patients (CDC) and Controls (Control). Non-significant results are not displayed.

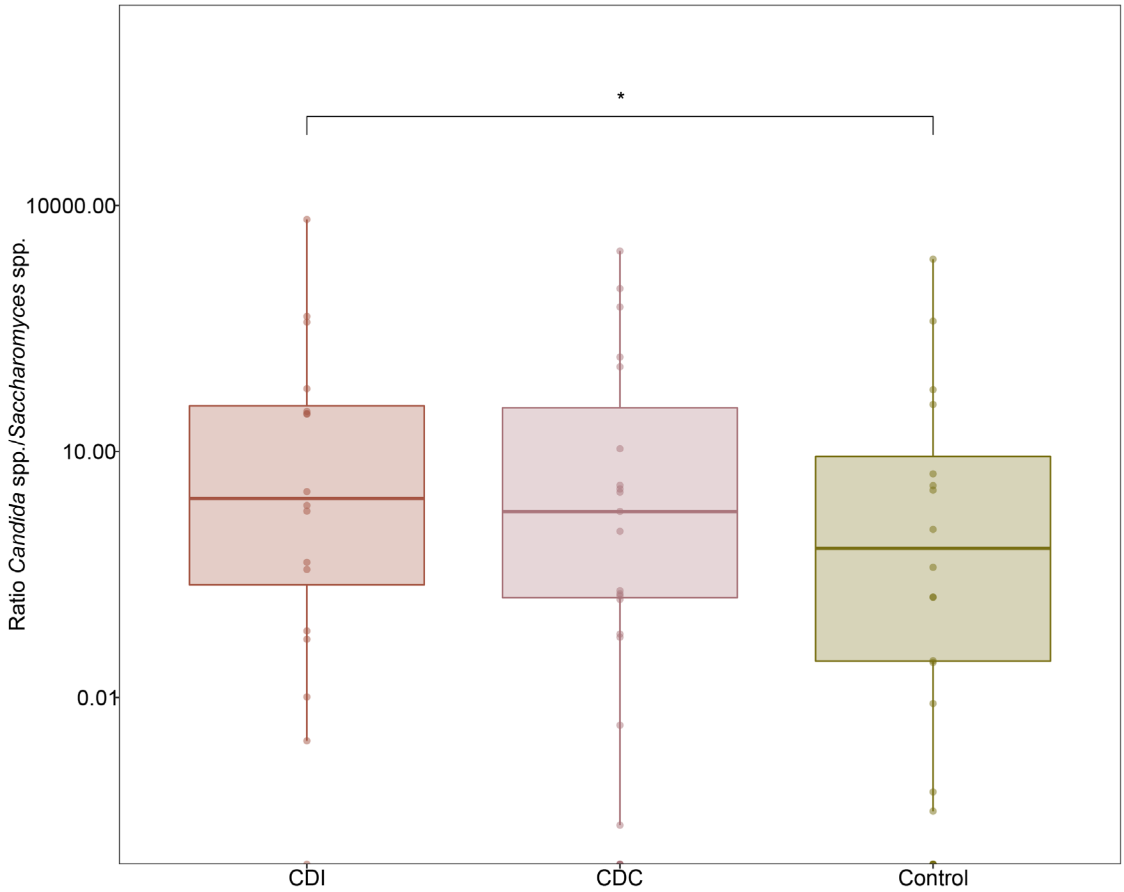


**Supplementary Figure 3.** Logarithmic scale of the ratio *Candida* spp. to *Saccharomyces* spp. In *C. difficile*-infected patients (CDI), asymptomatic *C. difficile*-colonised patients (CDC) and Controls (Control). $^{*}P \leq 0.05$ (Wilcoxon Rank Sum Test). Non-significant results are not displayed.

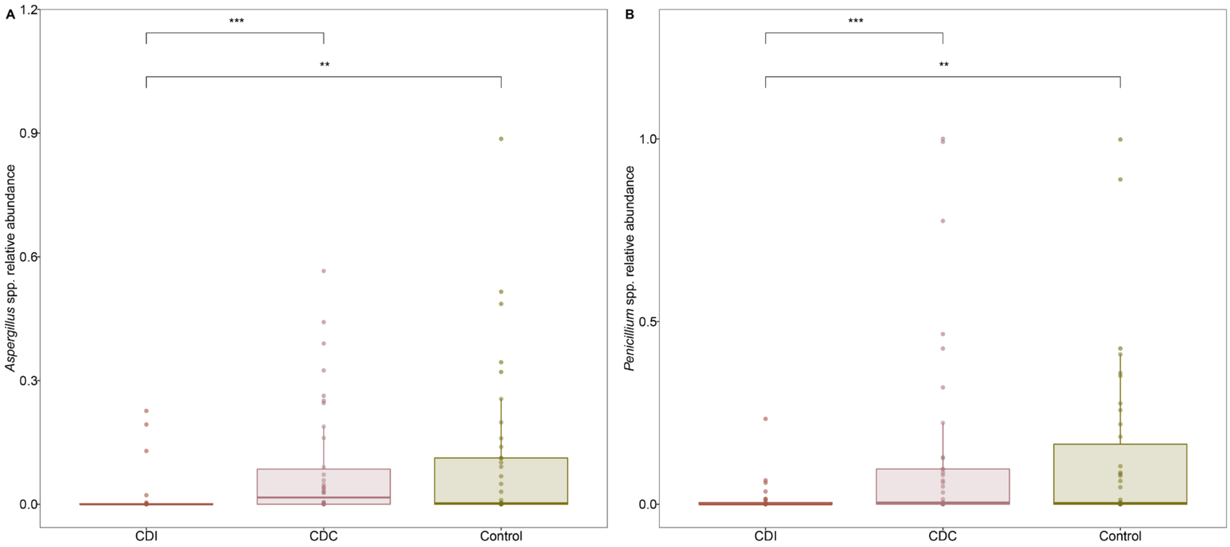


**Supplementary Figure 4.** Relative abundance of *Aspergillus* spp. (A) and *Penicillium* spp. (B) in patients with *C. difficile* infection (CDI), asymptomatic *C. difficile* colonised patients (CDC) and Controls (Control). Individual data points are displayed as circles. $^{***}P \leq 0.001$, $^{**}P \leq 0.01$ (Wilcoxon Rank Sum Test).

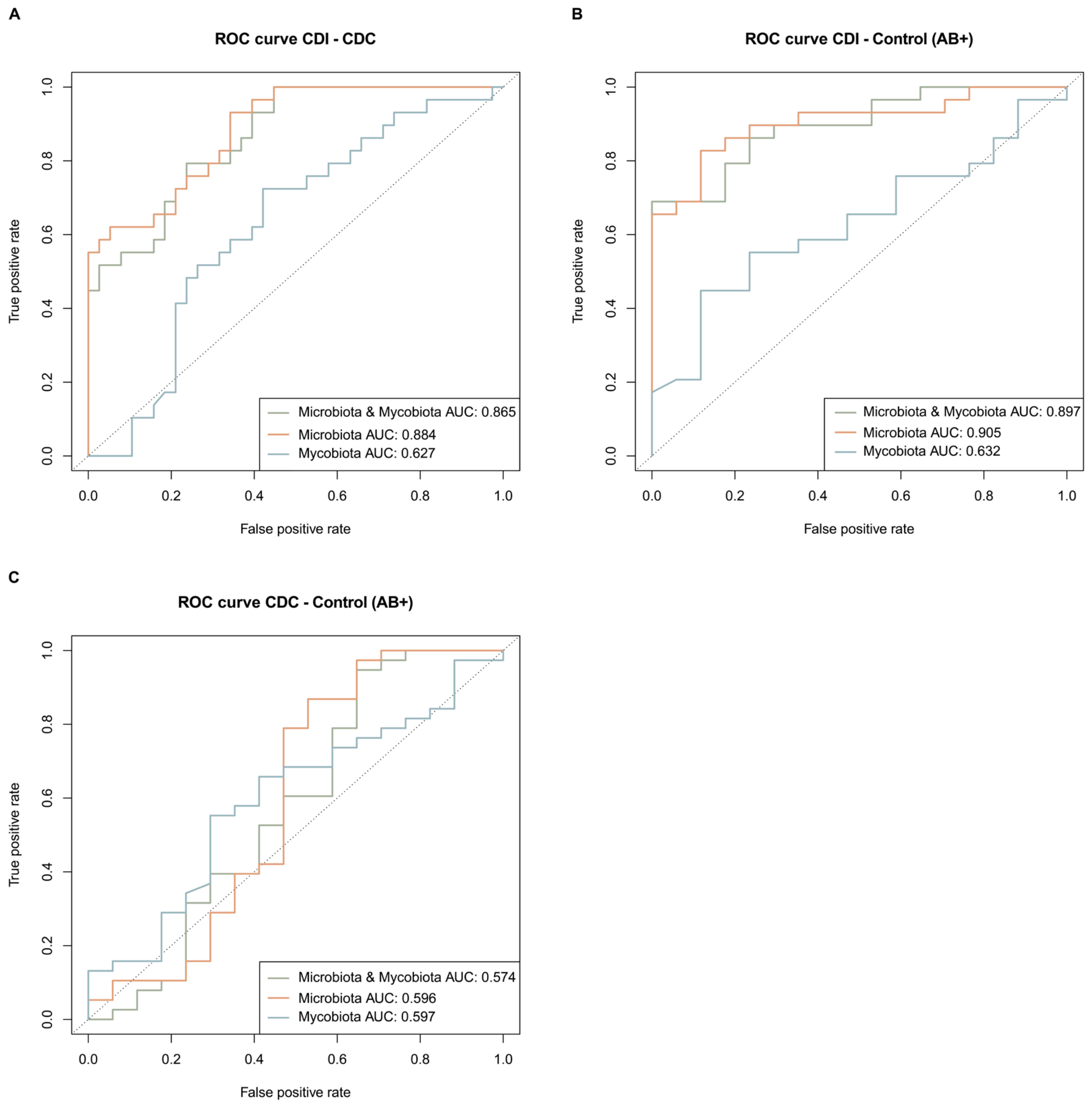


**Supplementary Figure 5.** Receiver Operating Characteristic (ROC) curve for the models distinguishing the groups. ROC curves are displayed for joint microbiota and mycobiota data (green), microbiota data (orange) and mycobiota data (blue) with corresponding Area Under the Curves (AUCs). A: models distinguishing *C. difficile*-infected patients (CDI) from asymptomatic *C. difficile*-colonised patients (CDC). B: models distinguishing *C. difficile*-infected patients (CDI) from Controls who received antibiotic treatment (AB+). C: models distinguishing asymptomatic *C. difficile*-colonised patients (CDC) from Controls who received antibiotic treatment (AB+).

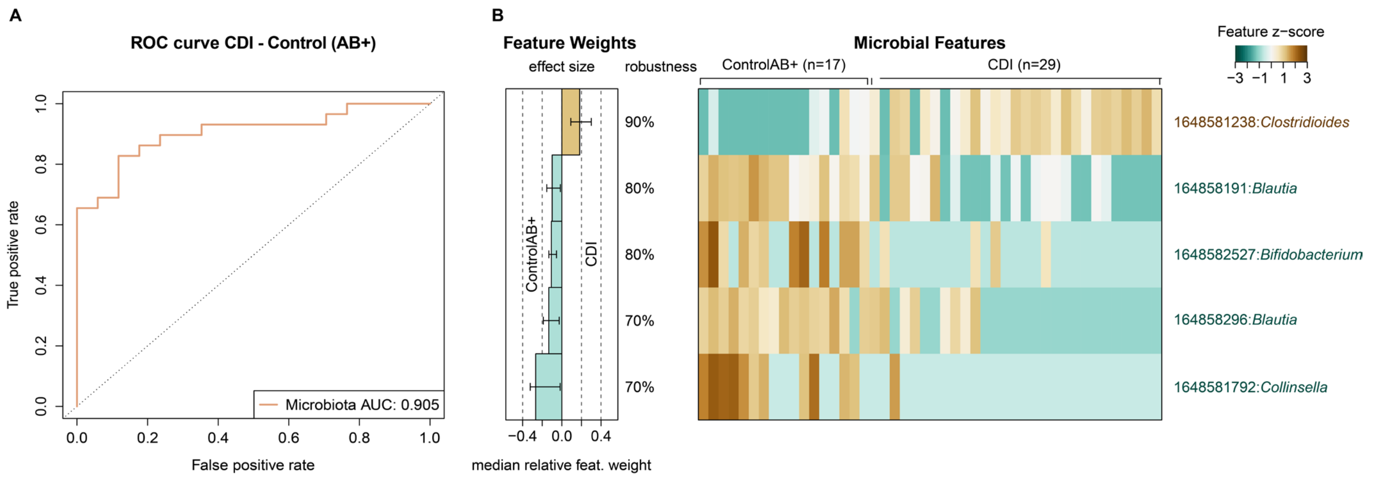


**Supplementary Figure 6.** Signature of gut bacterial OTUs associated to *C. difficile* infection (CDI) and Controls who received antibiotic treatment (ControlAB+). A: Cross-validation accuracy of the microbiota classifier is shown in the Receiver Operating Characteristic (ROC) curve with the mean Area Under the Curve (AUC). B: Median relative feature weights of the selected gut bacterial OTUs show the contribution of each marker OTU to the classification. Robustness of the selected features indicates the fraction of models containing the specific feature. The normalised values of the selected features across CDI and ControlAB+ is shown in the heatmap.

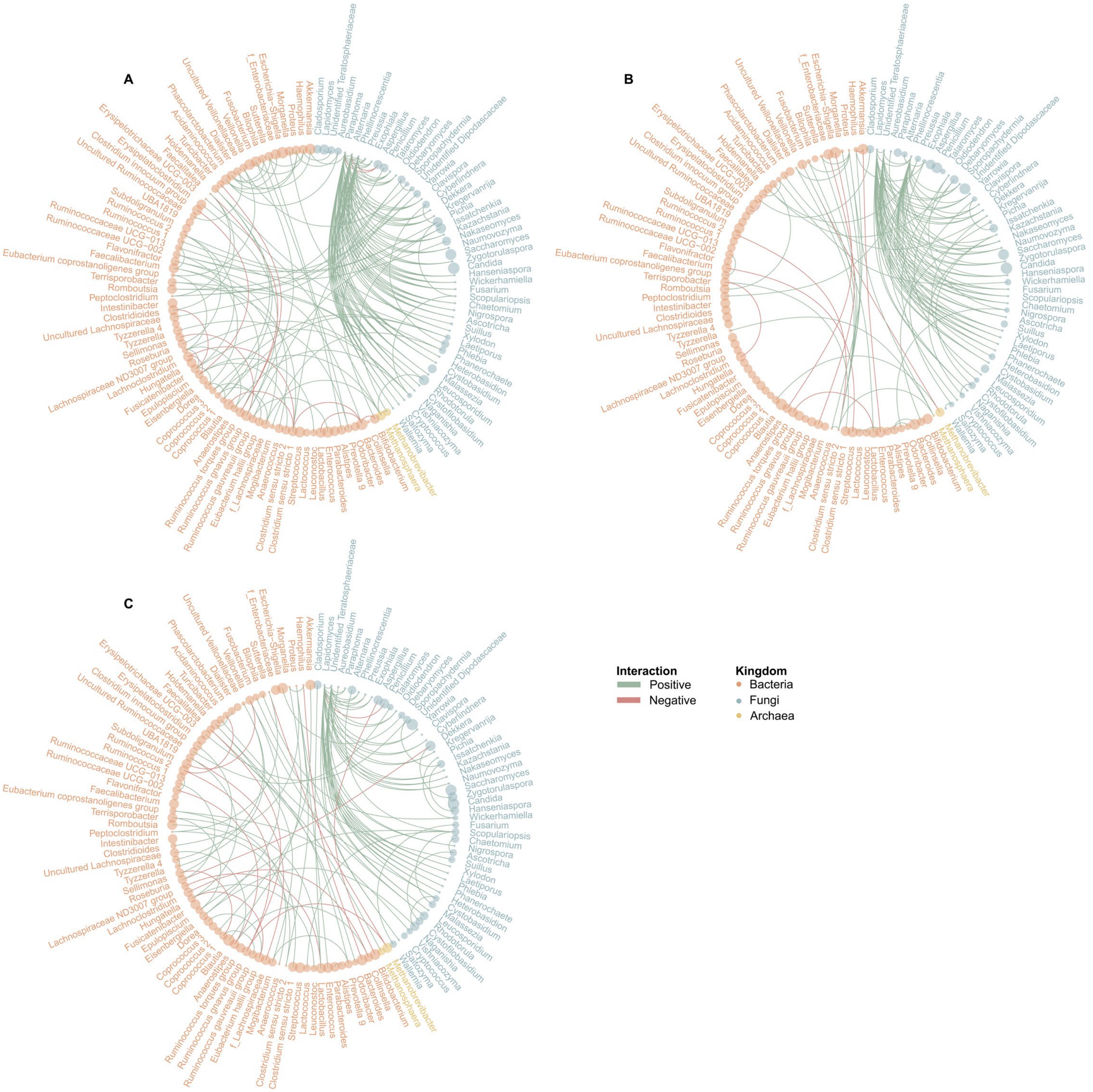


**Supplementary Figure 7.** Fungal-bacterial networks. Positive interactions are highlighted in green, while negative interactions are highlighted in red. Kingdoms are represented by colour; mean relative abundance of the genus are represented by point size. A: *C. difficile*-infected patients (CDI). B: *C. difficile*-colonised patients (CDC). C: Controls who received antibiotic treatment (ControlAB+).

[18] Wirbel J, Zych K, Essex M, Karcher N, Kartal E, Salazar G, et al. Microbiome meta-analysis and

cross-disease comparison enabled by the SIAMCAT machine learning toolbox. Genome Biology 2021 22:1 2021;22:1–27. https://doi.org/10.1186/S13059-021-02306-1.

[19] Kurtz ZD, Müller CL, Miraldi ER, Littman DR, Blaser MJ, Bonneau RA. Sparse and Compositionally Robust Inference of Microbial Ecological Networks. PLoS Comput Biol 2015;11:e1004226. https://doi.org/10.1371/journal.pcbi.1004226.

[20] Pedersen TL. ggraph: An Implementation of Grammar of Graphics for Graphs and Networks 2021.
